# KcatNet: A Geometric Deep Learning Framework for Genome-Wide Prediction of Enzyme Catalytic Efficiency

**DOI:** 10.1101/2025.03.09.642294

**Authors:** Tong Pan, Xin Cui, Huan Yee Koh, Yue Bi, Xiaoyu Wang, Yumeng Zhang, Shantong Hu, Geoffrey I. Webb, Lukasz Kurgan, Guimin Zhang, Jiangning Song

## Abstract

Enzyme turnover numbers (*K*_*cat*_) are fundamental kinetic constants that quantify enzymatic efficiency. Systematic studies of *K*_*cat*_ are essential for characterizing the mechanisms underlying proteomic composition and cellular metabolism. However, experimental measurements of *K*_*cat*_ remain limited and prone to noise. To address this, we present KcatNet, a geometric deep learning model designed for high-throughput prediction of *K*_*cat*_ in metabolic enzymes across all organisms, leveraging paired enzyme sequence and substrate representations. KcatNet consistently outperforms existing predictors, particularly for enzymes with high catalytic efficiency, and demonstrates strong generalization to enzymes that are dissimilar to those in the training set. Furthermore, KcatNet uncovers structural mechanisms and interaction patterns within enzyme-substrate complexes, providing insights into architectural principles that are inaccessible with existing methods by harnessing the representational power of large-scale protein language models. We applied KcatNet to genome-scale *K*_*cat*_ prediction across diverse yeast species, improving proteome allocation predictions by integrating its outputs into metabolic models. Experimental validation further confirmed the model’s ability to identify enzyme mutants with enhanced activity. By bridging the gap between sequence, structure, and function, KcatNet establishes a robust foundation for advancing understanding of molecular-level mechanisms and accelerating enzyme engineering efforts.

## Background

The enzyme turnover number, *K*_*cat*_, quantifies the number of substrate molecules converted to product per enzyme molecule per unit of time under substrate-saturated conditions [1]. As a fundamental parameter of catalytic efficiency, *K*_*cat*_ provides critical insights into cellular metabolism and enzyme evolution [2-4]. Incorporating organism-specific *K*_*cat*_ values into genome-scale metabolic models (GEMs) has advanced our understanding of protein allocation and cellular growth by accounting for the protein cost associated with specific reaction fluxes [5, 6]. Furthermore, the diverse catalytic efficiencies of enzymes are shaped by evolutionary and biophysical constraints imposed by natural selection [7, 8]. Therefore, investigating the differences in catalytic turnover rates underlying complex enzymatic mechanisms is essential for exploring biological systems from an evolutionary perspective [9]. Comparative analyses of catalytic activities across diverse enzymes and their mutants offer valuable insights into enzyme evolution, informing strategies for metabolic engineering [10].

*K*_*cat*_ is typically determined through experimental measurements. However, obtaining a complete *K*_*cat*_-ome for an organism via *in vitro* characterization faces significant challenges, including difficulty in purifying specific enzymes, limited substrate availability, and incomplete knowledge of required cofactors [11]. Currently, there are no high-throughput methods for experimentally measuring *K*_*cat*_ values and the evaluation process remains time-consuming and labor-intensive [12]. As a result, the number of annotated *K*_*cat*_ values for enzymes of individual species available in public databases such as BRENDA [13] and SABIO-RK [14] is limited and far below proteome-scale. For example, even in *Escherichia coli*, one of the most extensively studied model organisms, *in vitro K*_*cat*_ values are available for only ∼12% of enzymatic reactions [15]. Additionally, experimental *K*_*cat*_ values can span several orders of magnitude and vary widely due to differences in assay conditions [16]. This variability, combined with the sparse annotation of *K*_*cat*_ values, highlights the urgent need for computational tools capable of reliably predicting these values for enzymes on a proteome-wide scale. Such methods would enable exploration of enzyme activities and accelerate progress in biomolecular design [17].

Recent advances in artificial intelligence (AI) have spurred the development of novel predictors of *K*_*cat*_ values [18]. Methods such as DLkcat [19], TurNuP [20], and UniKP [21] leverage enzyme sequences and substrate features to predict *K*_*cat*_ values across diverse organisms, employing deep learning models or machine learning classifiers like gradient boosting [22] and extra trees [23]. Despite these advancements, these approaches face challenges in model training [24], limiting their ability to generalize effectively and accurately predict high *K*_*cat*_ values, which are underrepresented in experimental datasets. While experimentally measured *K*_*cat*_ data encompass a diverse range of enzyme sequences and substrates, they often lack detailed structural information about enzyme-substrate interactions. Manual efforts are typically required to correlate structural data with catalytic efficiency to uncover the structural mechanisms driving catalytic activity [25]. Predicting *K*_*cat*_ values using structural data remains a significant challenge [26], despite progress for specific organisms like *Escherichia coli*. For instance, Heckmann et al. [15] proposed a *K*_*cat*_ predictor that incorporates manually curated features, including catalytic site structures and reaction fluxes derived from parsimonious flux balance analysis. Although this approach demonstrates the potential of integrating enzyme structural data for *K*_*cat*_ prediction, its applicability is limited to a small subset of enzymatic reactions in well-characterized organisms like *E. coli*. Nevertheless, this remains a promising research direction, especially given recent breakthroughs in protein structure prediction, which have significantly expanded the availability of high-quality putative structural data for enzymes [27].

Motivated by the lack of broadly applicable and efficient methods that leverage structural data for *K*_*cat*_ prediction, we introduce KcatNet, an innovative tool based on a geometric deep learning model. KcatNet effectively captures structural information using paired enzyme sequences and substrate SMILES representations as input, enabling rapid and accurate *K*_*cat*_ prediction at the genomic scale. We demonstrate that KcatNet can capture the geometric contexts of enzyme-substrate interactions, even for enzymes without experimentally resolved structures, by working in conjunction with structure prediction methods. KcatNet outperforms existing methods in predicting *in vitro K*_*cat*_ values, exhibiting superior generalizability and accuracy, particularly for high *K*_*cat*_ values, while remaining agnostic to protein families. Additionally, KcatNet povideds interpretability by generating importance scores for residues involved in catalytic activity. It identifies spatially contiguous residue clusters proximal to or within the catalytic pocket, potentially linking these residues to functional roles beyond direct catalysis. We applied KcatNet to predict *K*_*cat*_ values across various yeast species, enabling the construction of the enzyme-constrained metabolic models that improve phenotype and proteome predictions. Finally, we used KcatNet to study the directed evolution of α-glucosidase. Among nine single-point mutations predicted by KcatNet to enhance catalytic efficiency, seven were experimentally validated to exhibit *K*_*cat*_ values exceeding those of the wild-type enzyme. Collectively, these results underscore KcatNet’s potential to advance enzyme engineering and metabolic modelling.

## Results

### Model overview

The KcatNet framework is illustrated in Fig. 1. KcatNet is a custom-designed deep learning model that predicts *K*_*cat*_ values at the genomic scale using enzyme-substrate pairs as inputs (Fig. 1a). The model’s design is grounded in the principle that enzymes catalyze reactions by binding substrate molecules in specialized regions known as catalytic pockets [28, 29]. Active residues within these pockets facilitate enzymatic reactions by recognizing and positioning substrates, contributing to both specificity and efficiency [30]. KcatNet is engineered to identify catalytic pockets and infer catalytic efficiency by analyzing enzyme substructures and uncovering associations between substrates and these pockets. The model comprises three main components, as shown in Fig. 1b: 1) an enzyme-substrate encoder module; 2) a graph-based residue-level partition module, and 3) and an interaction module that captures enzyme-substrate interaction patterns.

**Fig. 1.**
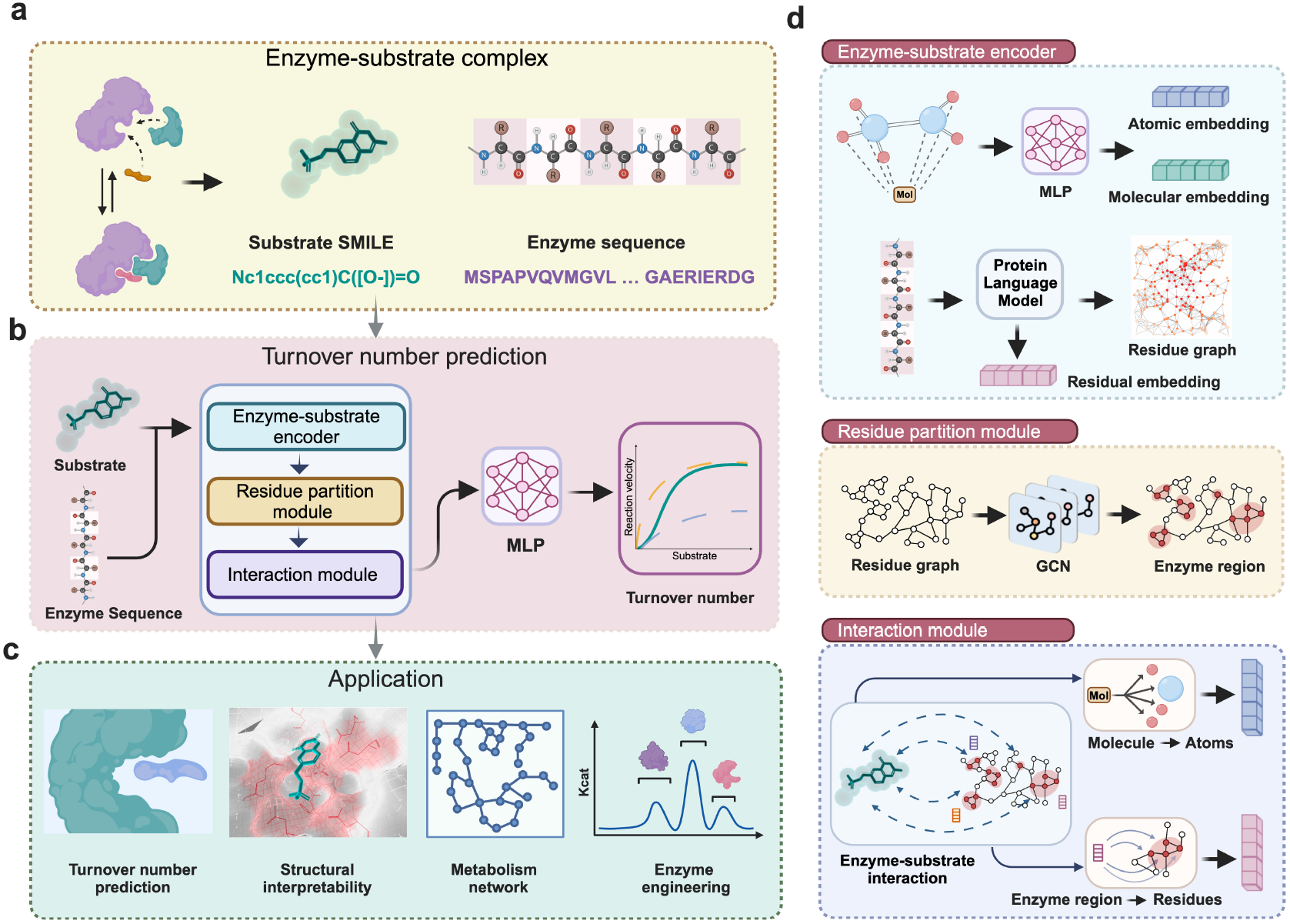
Overview of KcatNet. **a** KcatNet input: KcatNet takes enzyme sequences paired with substrate SMILES representations as inputs. **b** Architecture schematic: High-level visualization of the KcatNet model architecture. **c** Application: Predicted *K*_*cat*_ values generated by KcatNet are applied to downstream analyses. **d** Detailed components: Enzyme-substrate encoder: Substrates are represented at two levels—atomic (using atom type and property embeddings) and molecular (processed via a pretrained SMILES transformer to extract molecular fingerprints). Enzymes are modeled as residue-level graphs, where residue features are derived from pretrained language models (ProtT5 and ESM2) and linked through ESM2-derived contact maps; Residue partition module: Neighboring residues within the enzyme graph are identified and organized into local regions using graph convolutional networks (GCNs); Interaction module: Molecular fingerprints and enzyme region embeddings are iteratively refined using attention networks to model enzyme-substrate interactions. Substrate fingerprints are decomposed into atomic features, while enzyme regions are broken down into updated residue embeddings. This process is repeated, refining both enzyme regions and molecular fingerprints to produce the final *K*_*cat*_ prediction.

In the enzyme-substrate encoder module (Fig. 1d), representations of the enzyme and substrate are combined with complementary information generated by modern large language models. Specifically, we extract residue-level features from the enzyme sequence using ProtT5 [31] and ESM2 [27], two widely used deep learning-based protein language models (PLMs) trained on billions of amino acid sequences. These features encapsulate information about individual enzyme residues and their relationships within the context of the entire enzyme sequence. We concatenate these residue-level features to produce a comprehensive enzyme-wise sequence representation. For substrate representation, we use the SMILES format, a standard representation for encoding molecular structures. Substrates are encoded at two distinct levels: (i) Atomic level: Each atom is initialized with type and property embeddings, and (ii) Molecular level: The entire molecule is processed using a pretrained SMILES transformer [32]. In addition to the PLM-based enzyme sequence representations and SMILES-based substrate encodings, we encode local enzyme substructures through the graph-based residue-level partition module (Fig. 1d). Here, the enzyme structure is modeled as a graph that captures the spatial arrangement of its amino acids. We employ a graph convolutional neural network (GCN) to process these enzyme structure graphs [33], refining residue-specific representations by propagating embeddings among spatially proximal residues. For enzymes lacking experimentally resolved structures, we use ESM2 to generate residue contact maps, which quantify spatial relationships between residues. These residues are then grouped into distinct clusters using graph partitioning [34], a method that decomposes the graph into smaller, strongly connected subgraphs. The resulting residue partitions, represented as weighted graphs, undergo max pooling operations to produce partition-wise representations.

We process the three complementary inputs, which represent the substrate structure and the enzyme structure and sequence, using a Multilayer Perceptron (MLP) network to generate *K*_*cat*_ predictions. In essence, this network captures spatially local interaction patterns between substrates and catalytic pockets within the graph-based enzyme structure, contextualized by the PLM-based enzyme sequence space (Fig. 1d). An attention mechanism evaluates the interaction propensity of substrates with different residue regions to identify catalytic pockets. During training, the model updates both the substrate and enzyme partition-wise representations based on their contributions to the enzyme-substrate reaction. Message-passing operations are applied to both substrates and enzyme substructures, where each enzyme residue receives messages from its corresponding enzyme substructure, and each substrate atom receives messages from the molecular-level embedding. Through iterative training updates relying on the message passing, the final enzyme and substrate embeddings accurately represent substrate-catalytic pocket interactions. Pooling operations are then used to project these embeddings into scalar probability values representing turnover number predictions.

### KcatNet performance for enzyme turnover number prediction

We evaluated the accuracy of KcatNet for *K*_*cat*_ predictions on a benchmark set, as outlined in the Methods section, enabling a comparative assessment of our model’s performance against existing methods. Predictions were assessed using several performance metrics, including Pearson’s correlation coefficient (PCC), coefficient of determination (R^2^), root mean square error (RMSE), and mean absolute error (MAE), by comparing predicted *K*_*cat*_ values with experimentally observed data (Refer to the “Evaluation metrics” section in the Supplementary File).

We compared *K*_*cat*_ values predicted by KcatNet with experimental measurements for the same enzyme-reaction pairs, as shown in Fig. 2a. The results demonstrate a strong correlation between KcatNet’s predictions and experimentally measured *K*_*cat*_ values, achieving a PCC of 0.84 on the test set (Fig. 2a) and 0.94 on the entire dataset (Supplementary Fig. S4), respectively. We further compared KcatNet’s predictive performance with two recently published tools, namely DLKcat [19] and UniKP [21]. To ensure a fair evaluation, we retrained all models and assessed their performance on the same training and test sets. As shown in Fig. 2b. KcatNet achieved a substantially lower RMSE of 0.78 on the test dataset, outperforming DLKcat (****, *P* = 4.35 × 10^−10^) and UniKP (***, *P* = 3.15 ×10^−4^). Notably, compared to the second-best method (UniKP), KcatNet improved R^2^ by 18% (R^2^ = 0.69, ***, *P* = 4.07×10^−4^). Additionally, KcatNet yielded a lower MAE of 0.55, outperforming both DLKcat (****, *P* = 2.39 × 10^−11^) and UniKP (****, *P* = 1.32 × 10^−7^). Pairwise *P*-values for these comparisons are provided in Supplementary Table S3. These findings indicate that KcatNet’s predictions surpass those of previously published methods. Furthermore, when categorizing enzymes by their metabolic roles, KcatNet predicted higher *K*_*cat*_ values for enzymes involved in primary central and energy metabolism compared to those in intermediary and secondary metabolism (Fig. 2c). This observation aligns with previous findings [35] and provides additional evidence of the model’s accuracy.

**Fig. 2.**
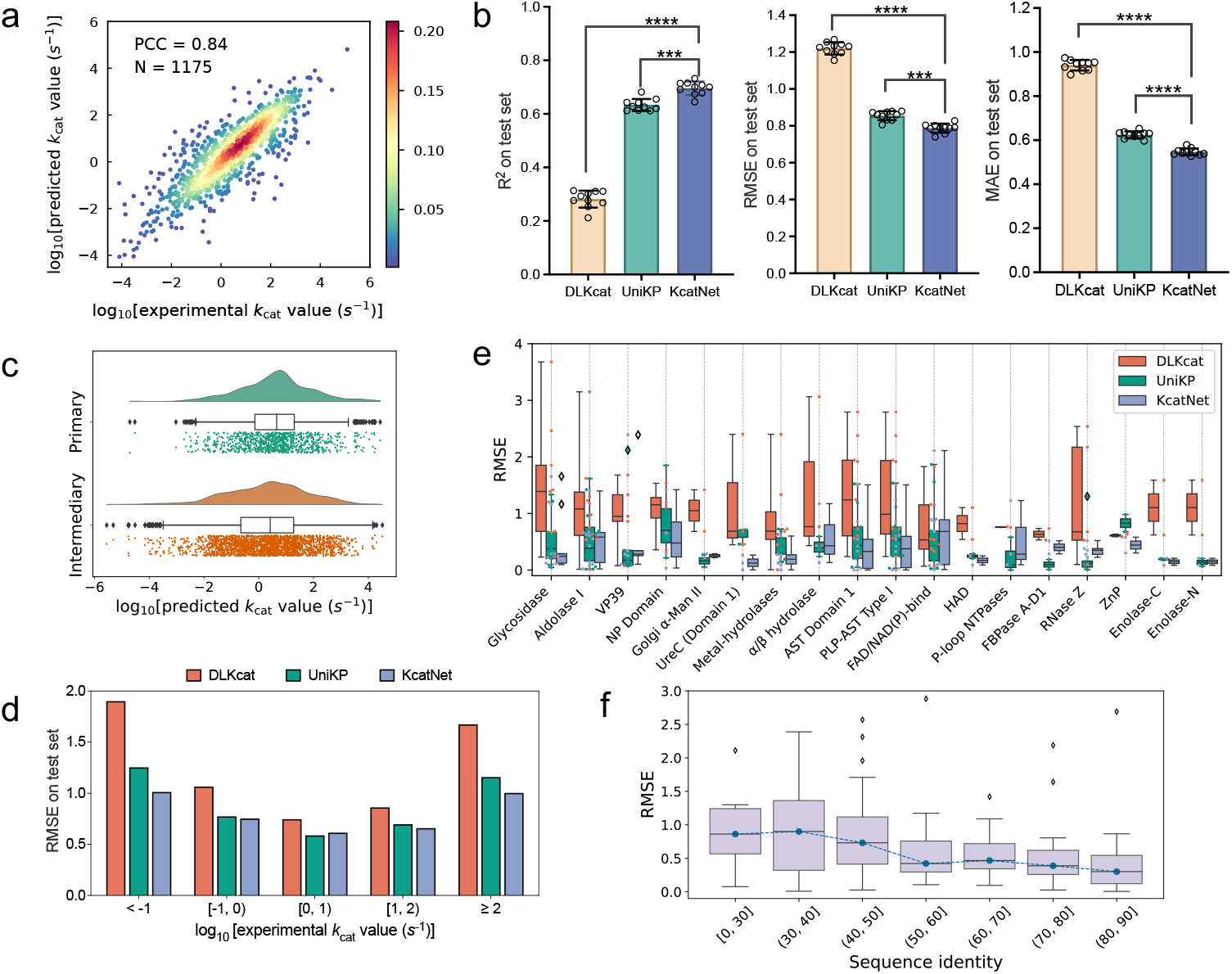
Predictive performance of KcatNet. **a** Scatter plot showing the Pearson coefficient correlation (PCC) between experimentally measured *K*_*cat*_ values and KcatNet-predicted *K*_*cat*_ values for the test dataset (*n* =1175). The color gradient represents the density of data points, ranging from blue (low density) to red (high density). **b** Performance comparison of KcatNet with baseline models DLKcat and UniKP in terms of the Root Mean Square Error (RMSE) (left) and Coefficient of Determination (R2) (right) between experimentally measured *K*_*cat*_ values and predicted *K*_*cat*_ values. Error bars represent the standard deviation of the mean based on 10 independent runs. Significance levels are indicated (two-sided *t*-test results: ∗∗∗, *P* ≤ 0.001; ∗∗∗∗, *P* ≤ 0.0001). **c** Distinct KcatNet predictions for enzymes in central energy metabolism versus intermediary and secondary metabolism. **d** Evaluation of KcatNet predictions for enzymes with varying *K*_*cat*_ values using RMSE scores on the test datasets. **e** Performance comparison of KcatNet and baseline models DLKcat and UniKP across various protein families based on RMSE. **f** RMSE for the test sets for KcatNet for different levels of maximal enzyme sequence identity compared to enzymes in the training set. For the box plots in **e** and **f**, the lower limit represents the lower quartile, the middle line represents the median, and the upper limit represents the upper quartile, respectively.

We observe that the distribution of *K*_*cat*_ values in the test dataset is skewed, with approximately 71% of values falling between 0.1 and 100, while only 14% exhibit high *K*_*cat*_ values (>100) (Extended Data Fig. 2a). Accurate identification of enzymes with high *K*_*cat*_ values is particularly important for enzymology and synthetic biology [5]. However, we find that existing models systematically overestimate low *K*_*cat*_ values and underestimate high ones (Supplementary Fig. S3, S5). To evaluate KcatNet’s performance in predicting high *K*_*cat*_ values, we stratified the analysis across different *K*_*cat*_ ranges (Fig. 2d). The results show that KcatNet provides accurate predictions, with RMSEs ranging from 0.61 to 1.01 across the entire spectrum of *K*_*cat*_ values, indicating predictions within one order of magnitude of experimental values. Notably, KcatNet consistently outperforms the two recently published methods, particularly excelling in predicting both very low and very high *K*_*cat*_ values. We attribute KcatNet’s superior performance in part to the enzyme representations derived from pretrained protein language models, as well as the model architecture that integrates structural features, as supported by our ablation analysis (Extended Data Fig. 3a, 3b). Furthermore, while the *K*_*cat*_ values in the test dataset originate from diverse organisms, we partitioned the test set into organism-specific subsets to evaluate KcatNet’s performance for individual organisms. We find that KcatNet delivers strong predictive performance across a wide variety of organisms, including those underrepresented in the training dataset, such as *Drosophila melanogaster* (Extended Data Fig. 3c, 3d).

### KcatNet captures enzyme homolog-independent patterns for *K*_*cat*_ prediction

To evaluate performance of KcatNet across protein families, we obtained protein family annotations from the CATH protein family database [36] and detailed the abundance of protein families in our test dataset in Supplementary Fig. S8. We found that KcatNet outperforms the two recently published methods across various protein families, including those with limited data, such as Golgi alpha-mannosidase II (Fig. 2e). This suggests that our model captures fundamental mechanisms of enzyme-substrate reactions rather than overfitting family-specific patterns.

Previous studies have shown that predictive performance is often influenced by the similarity between target enzyme sequences and those in the training dataset, as similar enzymes tend to share similar functions [1]. As expected, KcatNet performs exceptionally well in predicting *K*_*cat*_ for test enzymes highly similar to those in the training set, achieving a PCC of 0.90 (Supplementary Fig. S6). To assess KcatNet’s generalization ability, we tested its accuracy on previously unseen enzymes—those excluded from the training set (Extended Data Fig. 2b). KcatNet maintained a strong correlation between the predicted and experimental *K*_*cat*_ values, with a PCC of 0.84 (*N* = 1116). In this setting, KcatNet outperformed existing methods (Extended Data Fig. 2c), achieving a lower RMSE of 0.90 compared to DLKcat (****, *P* = 7.98 ×10^−5^) and UniKP (*, *P* = 2.13 ×10^−2^). Additionally, it achieved a higher R^2^ of 0.57, outperforming DLKcat (***, *P* = 2.07 ×10^−4^) and UniKP(*, *P* = 2.25 ×10^−2^), confirming the statistical significance of KcatNet’s superior performance. Pairwise *P*-values for these comparisons are provided in Supplementary Table S4. To further mitigate potential overfitting caused by similar enzyme entries, we evaluated the model’s robustness on enzymes dissimilar with those in the training dataset. As shown in Fig. 2f, we divided the test set into subsets based on their maximal enzyme sequence identity compared to the training set and calculated model performance at different levels of sequence identity. KcatNet generated accurate predictions even for test enzymes with low sequence identity to those in the training set.

### KcatNet identifies active sites involved in catalysis

A key strength of KcatNet lies in its ability to leverage enzyme structural features to provide interpretable context for catalytic efficiency predictions. By utilizing residue graphs, KcatNet is capable of identifying active sites within enzyme-substrate complexes that drive catalytic reactions, even without the prior knowledge of substrate-binding information.

We demonstrate that KcatNet can pinpoint spatially contiguous regions of residues proximal to or within the catalytic pocket that play critical roles in catalysis. As shown in Fig. 3a, using an enolase from *Pelagibaca bermudensis* HTCC2601 (PDB: 4H2H, chain A) as an example [37], we investigated the enzyme’s catalytic efficiency in catalyzing L-4-Hydroxyproline betaine. We visualized these predictions at the residue level by calculating the model’s activation scores (normalized attention weights) and mapping them onto the enzyme’s three-dimensional structure, with binding and active sites highlighted in red as retrieved from UniProt. KcatNet accurately identified all catalytic residues directly involved in the reaction by assigning them high activation scores. Additionally, all residues assigned high importance scores, though located at distinct sequence positions, were spatially proximal to the catalytic residues (Supplementary Fig. S9).

**Fig. 3.**
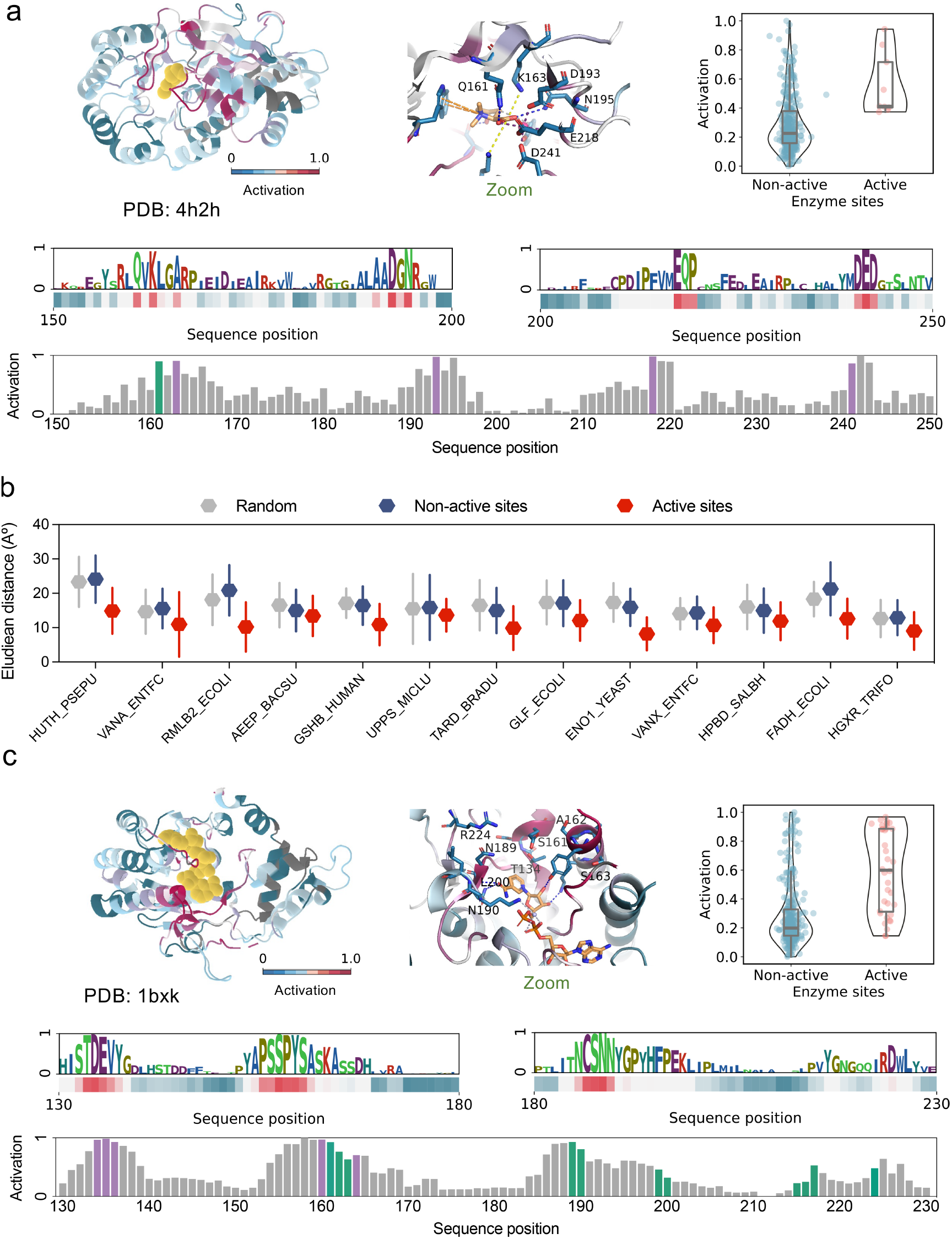
Analysis and interpretation of putative *K*_*cat*_ values generated by KcatNet. **a** Examples of important residues identified by KcatNet, showing consistency with enzyme functional sites for enolase (PDB ID: 4H2H, Chain A). The normalized attention weights (activation scores) are mapped onto the enzyme’s three-dimensional structure (upper panel), highlighting important residues in red and non-important residues in green. The substrate is represented in yellow. Key regions are magnified to display their detailed cartoon representation. Activation scores for both active and non-active sites are calculated, showing that important sites are typically assigned high activation scores (upper right). The sequence logo of residue importance scores is plotted (medium panel), where the color gradient represents the scores, ranging from blue (low activation scores) to red (high activation scores). Activation scores across the enzyme sequence are plotted (lower panel), with catalytic residues highlighted in purple, substrate binding sites marked in green, and the remaining sequence positions shown in gray. **b** Important residues identified by KcatNet with high activation scores are generally located close to enzyme catalytic sites. For each protein, we show the distribution of distances between the identified active sites (with high activation scores) and non-active sites (with low activation scores) and the nearest active site residue (from M-CSA). **c** Examples of important residues identified by KcatNet, showing consistency with enzyme functional sites for dTDP-glucose 4,6-dehydratase from *E. coli* (PDB ID: 1bxk, Chain A). Similar to **a**, activation scores across the enzyme sequence are plotted (lower panel), with catalytic residues highlighted in purple and substrate binding sites marked in green.

To quantify the spatial distribution of key residues identified by KcatNet as crucial for enzyme-substrate catalytic reaction efficiency, we analyzed 11 enzymes from our test dataset with detailed catalytic residue annotations in the MCSA database [38]. As shown in Fig. 3b, for each enzyme, we calculated the spatial distance between annotated catalytic residues and the active residues deemed important for turnover number prediction— those assigned high activation scores by KcatNet—as well as non-active residues (low activation scores). Our analysis revealed a distinct spatial distribution between active and non-active residues identified by KcatNet, with a significant portion of critical residues located near catalytic sites (Supplementary Fig. S9).

We hypothesize that some residues near or within the catalytic pocket play functional roles beyond direct catalysis. In some cases, the model identifies a small subset of residues with high activation scores that are involved in substrate binding. For example, as shown in Fig. 3c, we examined dTDP-glucose 4,6-dehydratase (PDB: 1BXK, chain A) from *E. coli* [39], which converts dTDP-glucose to dTDP-4-keto-6-deoxyglucose. The model identified a small group of residues with high activation scores that, while not directly participating in catalysis, are involved in substrate binding. Collectively, these results suggest that KcatNet effectively identifies regions near substrate binding and catalytic sites by calculating the functional importance of residues in predicting enzyme-substrate reaction efficiency, offering valuable insights into the structural basis of enzyme-substrate interactions.

### KcatNet accurately predicts *K*_*cat*_ values for mutant and wild-type enzymes

Next, we evaluated KcatNet’s ability to elucidate the effects of missense variants on *K*_*cat*_ values, a critical aspect of protein engineering. We separated catalytic reaction entries into wild-type and mutant enzymes and independently assessed the model’s predictions for each group. We observed a strong correlation between KcatNet’s predicted and experimentally measured *K*_*cat*_ values for both mutant enzymes (PCC = 0.85 for the test set, Fig. 4a; PCC = 0.94 for the entire dataset, Supplementary Fig. S10) and wild-type enzymes (PCC = 0.78 for the test set, Fig. 4b; PCC = 0.94 for the entire dataset, Supplementary Fig. S10). Compared to the two recently published methods, DLKcat and UniKP, KcatNet achieved lower RMSEs for both mutant enzymes (RMSE = 0.78 for the test set, Fig. 4c; RMSE = 0.52 for the entire dataset, Supplementary Fig. S11) and wild-type enzymes (RMSE = 0.77 for the test set, Fig. 4c; RMSE = 0.40 for the entire dataset, Supplementary Fig. S11). Similarly, KcatNet achieved higher R^2^ values between predicted and experimentally measured *K*_*cat*_ values for mutant enzymes (R^2^ = 0.71 for the test set, Fig. 4c; and R^2^ = 0.88 for the entire dataset, Supplementary Fig. S12) and wild-type enzymes (R^2^ = 0.59 for the test set, Fig. 4c; and R^2^ = 0.89 for the entire dataset, Supplementary Fig. S12), outperforming both DLKcat and UniKP.

**Fig. 4.**
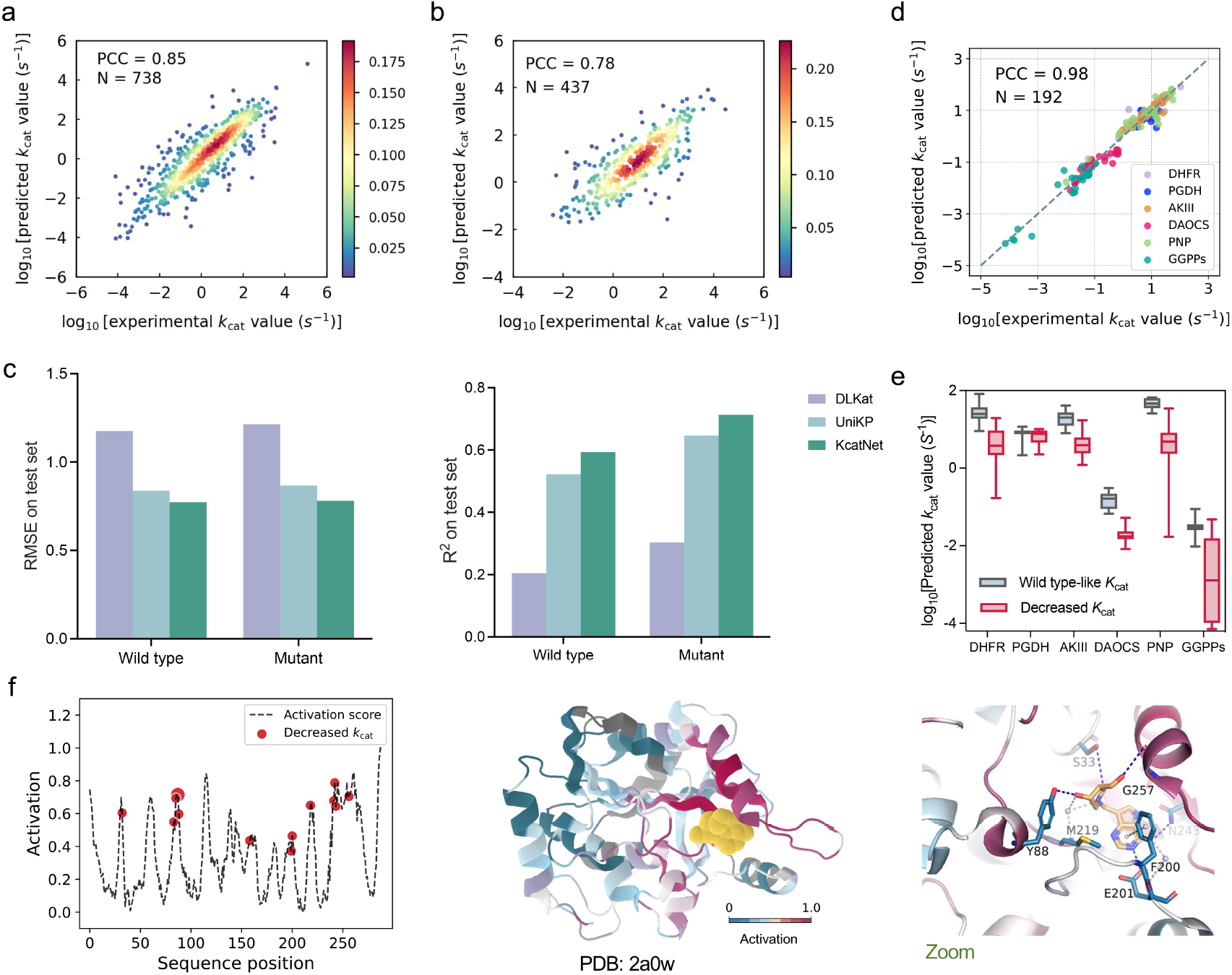
KcatNet discriminates *K*_*cat*_ values of enzymes and their mutants. **a-b** Scatter plots illustrating the Pearson coefficient correlation (PCC) between experimentally measured and predicted *K*_*cat*_ values for mutant enzymes (**a**) (*N*= 738) and wild-type enzymes (**b**) (*N*= 437). **c** Comparison of KcatNet with baseline models DLKcat and UniKP based on the Root Mean Square Error (RMSE) (left), and Coefficient of Determination (R2) (right) between experimentally measured and predicted *K*_*cat*_ values for wild-type and mutant enzymes. **d** Comparison between predicted and measured *K*_*cat*_ values for several well-studied enzyme– substrate pairs with extensive experimental mutagenesis data. Enzyme abbreviations: DHFR, dihydrofolate reductase; PGDH, d-3-phosphoglycerate dehydrogenase; AKIII, aspartokinase III; DAOCS, deacetoxycephalosporin C synthase; PNP, purine nucleoside phosphorylase; GGPPs, geranylgeranyl pyrophosphate synthase. Substrate abbreviations: G3P, glycerate 3-phosphate; L-Asp, l-aspartate; IPP, isopentenyl Diphosphate. **e** Comparison of predicted *K*_*cat*_ values for several mutated enzyme–substrate pairs between enzymes with wild-type-like *K*_*cat*_ and decreased *K*_*cat*_. **f** Normalized attention weights (activation scores) of wild-type PNP enzyme (PDB ID: 2A0W, Chain A) using inosine as the substrate (left). Red dots correspond to mutated residues with decreased *K*_*cat*_, marked on the curve according to their residue positions. Dot size indicates the number of mutant enzymes with mutations at that residue. Activation scores of residues predicted by KcatNet are mapped onto the enzyme structure (middle). Residues with high activation scores are highlighted in red (important), while others are shown in green (non-important). The substrate is depicted in purple. Important regions are magnified to show the cartoon representation (right).

Fig. 4d presents a subset of *K*_*cat*_ values for six mutant enzymes, each with at least 25 unique single or multiple amino acid substitutions. KcatNet accurately predicted *K*_*cat*_ values for these enzymes across 192 mutation entries, achieving a high PCC of 0.98. We further categorized the mutation data into wild-type and “reducing” mutations, defining the latter as those with *K*_*cat*_ values less than 50% of the wild-type enzyme. Fig. 4e illustrates that KcatNet effectively distinguishes between wild-type enzymes and those with reducing mutations, with wild-type enzymes exhibiting higher *K*_*cat*_ values. We also demonstrated that the residual-level importance scores identified by KcatNet, which reflect the extent to which residues influence catalytic efficiency through roles such as catalysis and substrate binding, are reliable and potentially biophysically significant. For example, by mapping the residue activation scores of the enzyme PNP (PDB: 2A0W, chain A) onto its three-dimensional structure [40], we observed a correlation between these activation scores and catalytic efficiency. Nearly all mutations at residues identified by KcatNet as highly important resulted in reduced *K*_*cat*_ values (Fig. 4f). This finding aligns well with the observation that functionally important residues in enzymes are typically conserved and that mutations in these residues often impair enzyme function [41].

### KcatNet’s predictions improve enzyme-constrained genome-scale metabolic models

Enzyme-constrained genome-scale metabolic models (ecGEMs) [42] simulate proteome allocation underlying key cellular properties such as growth rate, and rely heavily on genome-wide *K*_*cat*_ values, as protein costs in metabolic processes are determined by an enzyme’s effective turnover rate. The *K*_*cat*_ values predicted by KcatNet can facilitate the reconstruction of ecGEMs, enabling the investigation of proteome allocation patterns [43]. Li et al. *[19]* reconstructed original-ecGEMs for all 343 yeast/fungi species by assigning *K*_*cat*_ values from other substrates or organisms and, when necessary, introducing wildcards in the EC number to account for missing data. The original-ecGEMs reconstruction pipeline relied heavily on accurate EC number annotations and the availability of experimentally measured *K*_*cat*_ values in databases, ultimately yielding *K*_*cat*_ values for around 40% of enzymes. In contrast, deep learning–predicted *K*_*cat*_ values based on enzyme sequences and substrate SMILES significantly improved coverage, providing *K*_*cat*_ values for about 80% of enzymes *[19]*. The remaining gaps primarily involved generic substrates lacking defined SMILES representations, such as phosphatidate and thioredoxin [44]. Beyond achieving higher coverage, we applied the same original-ecGEMs pipeline but parameterized it with KcatNet-predicted *K*_*cat*_ values to assess their impact on proteome prediction accuracy. As shown in Fig. 5a, integrating KcatNet-predicted *K*_*cat*_ values into ecGEMs improved performance over the original ecGEMs, yielding more accurate growth rate predictions for four yeast species across diverse carbon sources and oxygen conditions. Compared to models parameterized with DLKcat-predicted *K*_*cat*_ values, our predictions reduced the mean squared error between the predicted growth rates and published quantitative data in 16 out of 22 environment-species combinations.

**Fig. 5.**
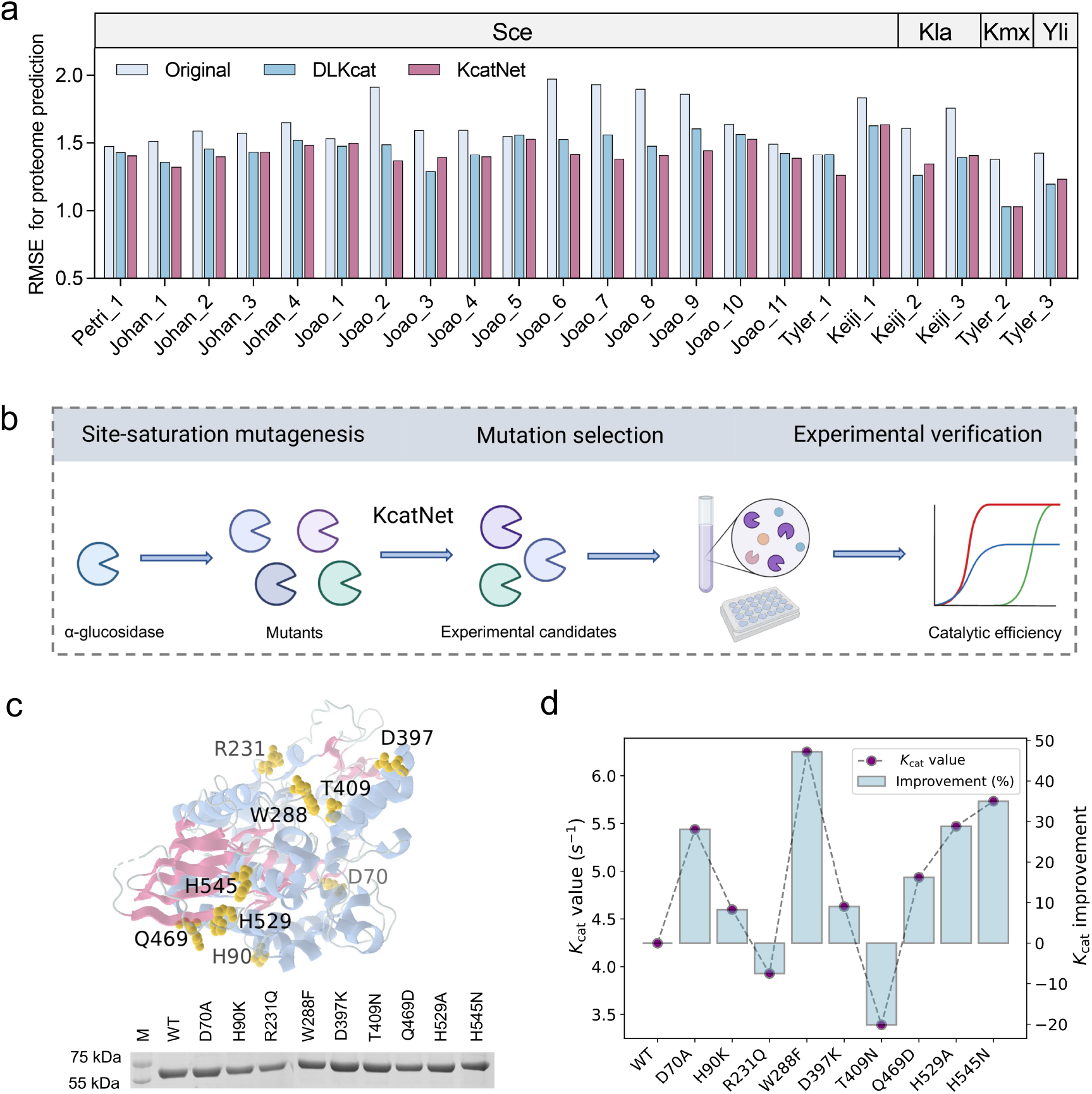
Application of KcatNet to ecGEMs and enzyme engineering. **a** Comparison of the original-ecGEM with ecGEMs parameterized using *K*_*cat*_ values from KcatNet and DLKcat for predicting quantitative proteome data, evaluated against experimental results using root mean square error (RMSE). Proteome predictions were generated for four yeast species (Sce, *Saccharomyces cerevisiae*; Kla, *Kluyveromyces lactis*; Kmx, *Kluyveromyces marxianus*; Yli, *Yarrowia lipolytica*) across distinct culture conditions. **b** KcatNet-based *in silico* screening of 10,545 α-glucosidase missense mutants to identify potential variants with improved *K*_*cat*_ for experimental validation. **c** Experimental impact of α-glucosidase missense variants: The nine mutated residues are plotted on the 3D structure, along with their protein abundance assessed by western blot (left) and experimentally measured *K*_*cat*_ values of the variants (right).

### Harnessing KcatNet for enzyme engineering

Enzyme engineering aims to develop more efficient enzyme variants through directed evolution. However, identifying effective evolutionary pathways is challenging, as it requires an understanding of molecular mechanisms, such as gene expression and protein folding, while navigating biological and physical constraints. We applied KcatNet in a prospective study to guide the directed evolution of α-glucosidase [45], a key enzyme in the amylolytic pathways of various organisms. α-Glucosidase catalyzes the hydrolysis of α-D-glucosides and facilitates transglucosylation, enzymatic activities extensively used in the synthesis of oligosaccharides and α-D-glucosides. Despite its importance, progress in designing α-glucosidase mutants with enhanced efficiency has been limited.

We employed KcatNet to prioritize α-glucosidase variants with potential improvements in catalytic efficiency (Fig. 5b). A comprehensive library of single-point mutations was computationally generated by substituting each residue in the α-glucosidase sequence with all possible alternative amino acids. These variants were then screened in silico using KcatNet to predict their corresponding *K*_*cat*_ values. Based on the predictions, nine residues were selected as candidates for experimental validation, each harboring multiple mutations predicted to improve *K*_*cat*_ (Fig. 5c). For each of these residues, one mutation was randomly chosen for downstream experimental testing. In addition, we assessed the quality of the selected variants using two established *in silico* tools: the sequence-based ESM-1v [46] and the structure-based ProteinMPNN [47], both of which were experimentally validated by Sean R. Johnson et al. [48] for their ability to predict the expression and folding of enzymes into soluble, functional forms. All variants achieved favorable scores, suggesting high potential for proper folding and functionality. Experimental results indicated that among the nine selected mutants, seven exhibited highe *K*_*cat*_ values than the wild-type enzyme (Fig. 5d). These results demonstrate that KcatNet could effectively aid in enzyme engineering by identifying promising variants for experimental evaluation.

## Discussion

A systematic and comprehensive investigation of enzyme efficiency is essential for unraveling their molecular catalytic mechanisms and evolutionary trajectories, enabling the design of enzymes to address current challenges in medicine, engineering, and planetary health [49]. However, predicting enzyme kinetic constants on a genome-wide scale remains a significant challenge due to variability in organismal fitness and the diversity of biochemical reactions (Supplementary Fig. S2). In this study, we introduce KcatNet, a sophisticated deep learning-based framework that accurately predicts the kinetic constant *K*_*cat*_ on a genome-wide scale. KcatNet processes enzyme sequences and substrate SMILES representations as inputs, consistently delivering accurate *K*_*cat*_ predictions by effectively capturing site-specific interaction patterns between substrates and enzyme active sites. Our model identifies spatially contiguous enzyme regions, including residues directly involved in catalysis and those critical for substrate binding, providing a detailed mapping of enzyme-substrate interactions and catalysis that is not accessible with existing approaches.

KcatNet generates accurate predictions of *in vitro K*_*cat*_ values for enzymes across a diverse collection of organisms, achieving PCCs > 0.80 (Extended Data Fig. 3d). We empirically demonstrate that KcatNet outperforms DLKcat, a leading deep learning model that incorporates substrate information and protein sequences, and UniKP, a recent tree-based ensemble model, in predicting *K*_*cat*_ for both mutant and wild-type enzymes. KcatNet also exhibits strong generalizability, accurately predicting *K*_*cat*_ for enzymes and reactions absent from the training set (Extended Data Fig. 2b, 2c). Even when evaluated on enzymes with low similarity to those in the training set, KcatNet maintains a low RMSE between measured and predicted *K*_*cat*_ values. Additionally, KcatNet effectively distinguishes catalytic efficiency differences for enzymes in various metabolic contexts, predicting higher turnover numbers for enzymes in high-flux pathways of primary central and energy metabolism compared to those involved in intermediary and secondary metabolism.

Identifying enzymes with high catalytic efficiency is a central goal in enzymology and synthetic biology [10]. KcatNet demonstrates superior performance in predicting high *K*_*cat*_ values, particularly in the context of highly imbalanced data where such high instances are rare. Traditional machine learning models have historically struggled with this imbalance, often failing to extract crucial information from limited high-*k*_*cat*_ data. By categorizing our dataset according to enzyme class (first-level EC numbers) and comparing the average measured *K*_*cat*_ values with the average predicted values for each class, we found that KcatNet consistently predicted higher *K*_*cat*_ values for faster EC classes and lower values for slower ones (Extended Data Fig. 4b). This result underscores KcatNet’s effectiveness in capturing the intrinsic catalytic properties of enzymes, as evidenced by its accurate predictions of *K*_*cat*_ values across different enzyme classes (Extended Data Fig. 4c).

We attribute KcatNet’s superior performance in part to the enzyme representations generated by pretrained large language models. Our analysis revealed that such representations play a critical role in predicting *K*_*cat*_ values (Extended Data Fig. 3a). Notably, even when predictions were based solely on enzyme information—without accounting for the catalyzed reaction—a respectable coefficient of determination was achieved. This finding suggests that the intrinsic properties of enzymes significantly influence turnover numbers, likely reflecting varying selection pressures on catalytic turnover driven by enzyme utilization. Interestingly, additional substrate data might yield only moderate performance improvements if the model struggles to capture key interactions essential for inferring reaction efficiency, particularly due to the absence of explicit structural enzyme-substrate binding information. The role of active sites in catalysis—whether through substrate binding, catalytic transformation, or direct participation in the reaction—has been extensively studied in structural and biochemical research [50]. However, existing models for predicting *K*_*cat*_ often overlook the detailed molecular interactions between enzyme active sites and substrates, largely because most enzyme-substrate complex structures are unavailable. KcatNet addresses this limitation by leveraging structural characterizations of enzymes to improve the interpretability of catalytic efficiency predictions. Our study demonstrates that KcatNet, when applied to multiple enzymes, effectively captures structurally significant site-specific interactions between enzymes and substrates. It uncovers spatially contiguous regions of residues near active sites linked to key functional roles, such as ligand-binding sites and catalytic residues. This approach offers valuable biological insights into enzyme-substrate interactions and catalysis, providing a detailed map of the underlying architecture—something that existing methods struggle to achieve, even without prior ligand-binding information.

Despite KcatNet delivering reliable *K*_*cat*_ predictions with performance surpassing existing methods, several challenges remain. One significant challenge is the high variance in *K*_*cat*_ measurements for the same enzyme-reaction pairs across different studies. Although we use a manually curated *K*_*cat*_ dataset for model construction and evaluation, this variance persists. It likely stems not only from database errors and differences in experimental procedures or assay conditions, such as temperature and pH. Another challenge arises with reactions involving multiple substrates or heteromeric enzyme complexes. KcatNet currently takes one substrate and one protein as input, while multiple different substrates or proteins may be involved in such reactions. This can result in multiple predicted *K*_*cat*_ values for a single reaction. In these cases, a more refined approach to predict a single *K*_*cat*_ value for multi-substrate or heteromeric enzymes would be ideal. Moreover, the current model does not account for environmental factors, a critical limitation when simulating real experimental conditions. *K*_*cat*_ values can vary considerably depending on factors like pH and temperature. Since experimental conditions are rarely documented in enzyme kinetic parameter databases, such information was unavailable as inputs for our prediction model. It is also important to note that other kinetic parameters, such as the Michaelis constant (*K*_*m*_), do not improve KcatNet’s performance in predicting *K*_*cat*_. Future work on developing methods to simultaneously predict *K*_*cat*_ and *K*_*m*_ values, aligning with experimentally determined *K*_*cat*_/*K*_*m*_ ratios, could enable comprehensive parameterization of enzyme kinetics and provide valuable insights into cellular physiology.

A key application of KcatNet is to facilitate the reconstruction of enzyme-constrained metabolic models (ecGEMs) for virtually any organism by predicting genome-scale *K*_*cat*_ profiles as kinetic parameters. We parameterized enzyme-constrained metabolic models using the predicted in vitro kinetic constants to align with existing experimental growth data for multiple yeast species. Our results demonstrate that KcatNet provides comprehensive *K*_*cat*_ predictions that significantly improve proteome allocation predictions. Additionally, KcatNet accurately captures changes in *K*_*cat*_ due to single amino acid substitutions, identifying mutations in catalytic residues that are likely to result in loss of enzyme function. The application of KcatNet to α-Glucosidase in directed evolution highlights its transformative potential for synthetic biology and biochemistry research. KcatNet effectively identified high-activity α-Glucosidase variants, with seven out of nine single-point mutations predicted to enhance catalytic efficiency, exhibiting *K*_*cat*_ values exceeding those of the wild-type enzyme. In conclusion, KcatNet can be broadly applied for monitoring biocatalysis, guiding enzyme engineering, and advancing our understanding of enzyme evolution and metabolic pathways.

## Conclusions

KcatNet represents a scalable and interpretable deep learning framework for the accurate prediction of enzyme turnover numbers (K<sub>cat</sub>) across genomic landscapes. By integrating enzyme sequence and substrate structure information through a geometric learning architecture, KcatNet robustly captures spatial determinants of catalysis and substrate binding, even in the absence of experimentally resolved enzyme structures. Beyond superior predictive performance, KcatNet offers mechanistic interpretability by highlighting functionally relevant residue clusters within catalytic regions. Applied to genome-scale metabolic models in diverse yeast species, KcatNet enhances predictions of proteome allocation and organismal phenotype. Moreover, its capacity to identify catalytically beneficial mutations demonstrates its utility in guiding enzyme engineering and synthetic biology. These findings establish KcatNet as a powerful tool for bridging sequence–structure–function relationships and accelerating the rational design of biocatalysts across biomedical and biotechnological domains.

## Methods

### Training and test datasets

We curated a dataset of *K*_*cat*_ measurements for both wild-type and mutant enzymes, which we used to construct and evaluate our predictive model. For the *K*_*cat*_ values of wild-type enzymes, we utilized the dataset curated by Kroll *et al*. This dataset derived the *K*_*cat*_ values from three databases—BRENDA, UniProt, and Sabio-RK—linking *K*_*cat*_ valus with corresponding enzyme sequences, reactant IDs, and reaction equations. To minimize noise in the data, such as errors from transcription or unit misplacements, more than half of the data points were manually verified to ensure *K*_*cat*_ values matched those reported in the original papers.

For the *K*_*cat*_ values of mutant enzymes, we collected data from the BRENDA (Release 2023.1) and SABIO-RK databases, focusing on entries with mutation annotations in their descriptions. Each dataset was linked to substrate details, organism information, UniProt IDs, and/or enzyme EC numbers. When multiple *K*_*cat*_ values were available for the same enzyme-substrate pair, we excluded values less than 1% of the maximum and calculated the geometric mean of the remaining values. Substrate SMILES, representing the substrate structures, were retrieved from the PubChem database by matching substrate names. Entries without corresponding SMILES were excluded. For protein sequences, we obtained entries with UniProt IDs directly from the UniProt database using their mapping service. For entries without UniProt IDs, sequences were retrieved from both the UniProt and BRENDA databases based on EC number and organism information. Entries returning multiple sequences were omitted.

Given that most *K*_*cat*_ values in the BRENDA and SABIO-RK databases lacked experimental condition details such as pH and temperature, we omitted this information from our curated dataset. As a result, we obtained 11,757 final entries for model training and testing purposes, including 11,288 unique reactions and 7,441 unique enzymes. All *K*_*cat*_ values were converted to a logarithmic scale. This dataset was randomly divided into an 80% training dataset for model construction and a 20% test dataset for performance evaluation and comparison.

### Enzyme sequence and structure embedding

The model takes as input an enzyme sequence paired with the SMILES representation of a molecular substrate and outputs the predicted *K*_*cat*_ value. We represent an enzyme sequence *S*= {r_1_, r_0_, …, r_*n*_}, consisting of *n* residues, as an attributed graph. This graph representation of the enzyme, denoted as *ENZ* = (*V, A, X*), embodies the three-dimensional structure of the input enzyme. In this representation, the residue set *V* ⊆ *S* servers as the graph’s nodes, the adjacency matrix *A* (of size *n* × *n*) quantifies the connectivity of these nodes based on their spatial distance, and the residue feature matrix *X* ∈ ℛ^*n*×*d*^ encapsulates the properties of the residues.

For an enzyme sequence S, we generate residue-level feature embeddings using two pretrained protein language models: ProtT5 and the model proposed by Bepler and Berger [46], resulting in embeddings 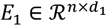 and 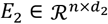, where *d*_1_ defaults to 1024 and *d*_2_ defaults to 1280, respectively. ProtT5 is a deep learning-based language model pre-trained on datasets comprising 393 billion amino acids. Bepler and Berger’s model is a bidirectional long short-term memory (Bi-LSTM) neural network trained on three types of data: 1) The protein’s SCOP classification based on its general structure; 2) The self-contact map of the protein’s structure, and 3) Sequence alignments of homologous proteins. These enzyme vectors encapsulate information about individual residues and broader protein-level characteristics. Additionally, we represent the enzyme’s spatial structure as an *n* × *n* residue adjacency matrix, generated by Bepler and Berger’s model, which encodes the 2D structure of the enzyme S.

### Substrate molecular embedding

Substrate molecules are represented using the SMILES format. For a substrate molecule *Sub*={r_1_,r_2_,…,r_m_} consisting of *m* atoms, the representation is embedded at two levels: the atomic level and the molecular level. At the atomic level, each atom is initialized with type and property embeddings, forming the atomic feature matrix *S*_atom_ ∈ **ℛ**^m×43^, which encapsulates the atom’s characteristics. Atom type features are derived from one of nine possible element groups: B, C, N, O, P, S, Se, halogens, and mMetals. The physicochemical properties include atom degree, the number of implicit hydrogens, atom hybridization, and chirality. At the molecular level, we obtain a comprehensive substrate representation *S*_mol_ ∈ **ℛ**^1×1024^ using a pretrained SMILES transformer, which captures the overall properties of the substrate.

### Overview of the KcatNet model

The KcatNet model leverages enzyme sequences and one of the substrates to accurately predict genome-scale enzyme turnover numbers. This is achieved through a structure-characterized deep learning framework that analyzes enzyme substructures and uncovers correlations between substrates and catalytic pockets to infer catalytic efficiency. This framework consists of three major components: 1) An enzyme-substrate representation encoder module embeds enzymes using a geometrically invariant representation and employs a multi-scale substrate representation at both the atomic and molecular levels, facilitating message propagation between these scales. 2) A graph-based residue-level partition module broadcasts embeddings among spatially proximal residues, grouping closely interacting residues into distinct partitions, and 3) A catalytic complex turnover number predictor captures the interaction patterns between enzyme residue partitions and their corresponding substrates, enhancing the prediction of enzyme kinetic turnover numbers.

### The enzyme-substrate representation encoder

We employed a graph encoder based on Principal Neighborhood Aggregation (PNA), a variant of graph convolutional network, to compute and propagate residue-level representations of enzymes. This encoder enhances message passing between spatially proximal residues within the enzyme structure by incorporating multiple aggregators and degree scalers, accounting for the characteristic residue density at protein functional sites.

Initially, we generate the enzyme embedding *E*_*ini*_ ∈ ℛ^*n*×*h*^ by concatenating and reducing the dimensionality of enzyme representations to a hidden dimension *h* using a multilayer perceptron (MLP) network, with *h* set to 200 by default:

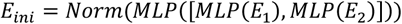

where Norm refers to layer normalization applied to the input features to stabilize the learning process, and the brackets [] indicate the concatenation operation. We use the Rectified Linear Unit (ReLU) activation function.

The initial enzyme representation *E*_*ini*_ ∈ ℛ^*n*×*h*^, along with the adjacency matrix *A* of size *n* × *n*, is fed into a PNA module. This module produces residue-level representations 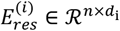 through message passing between spatially proximal residues, where *d*_*i*_ denotes the embedding dimension of the *i*th convolutional layer:

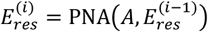

For the substrate, we generate the embedding *S*_*ini*_ ∈ ℛ^1×*h*^ by summing individual atom-level embeddings *X*_*atom*_ ∈ ℛ^m×43^ into a single ligand representation *X*_*lig*_ ∈ ℛ^1×*h*^ based on the atoms’ importance scores and concatenating global features of the molecular compound *X*_*mol*_ ∈ ℛ^1×1024^ using an MLP network, with *h* set to 200 by default:

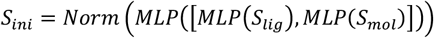

We aggregate ligand atoms into a unified ligand representation *S*_*lig*_ ∈ ℛ^1×*h*^ by summing atom embeddings 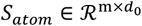 according to their importance scores:

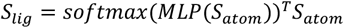

### The graph-based residue-level partition module

The graph-based residue-level partition module groups residues into several partitions based on their spatial proximity. For a given graph defined by the adjacency matrix *A* ∈ {0,1}^*n*×*n*^ and the residue-level feature matrix *E*_*res*_ ∈ ℛ^*n*×*h*^, our model applied a graph convolutional network (GCN) to produce an enzyme partition assignment matrix *A*_*region*_ ∈ ℛ^k×*n*^, where *k* represents the number of enzyme regions for the *i*th convolutional layer. The enzyme partition features *E*_*region*_ ∈ ℛ^k×h^ is subsequently derived by pooling residue features according to their assigned regions.

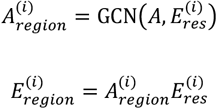

### The catalytic complex turnover number predictor

The catalytic complex turnover number predictor consistently estimates *K*_*cat*_ values by modeling the local interaction patterns between substrates and enzyme pockets. Using the substrate molecular representation *S* ∈ ℛ^1×*h*^ and the enzyme partition features *E*_*region*_ ∈ ℛ^k×h^ as inputs, the model infers the propensity of specific enzyme regions for enzyme-substrate interactions by calculating the relationship between the substrate molecule and various enzyme partitions through an attention mechanism. The substrate molecular representation and enzyme partition representations are then updated as follows:

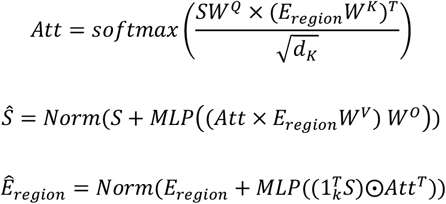

where *W*^Q^, *W*^K^, *W*^V^, and *W*° are learnable parameters, *d*_*K*_ is a constant scalar factor, and *Norm* refers to layer normalization.

A message-passing operation is then applied to independently facilitate information exchange within the substrate and enzyme substructures. Specifically, substrate representations at both the atomic and global molecular levels, as well as enzyme representations at the residue and partition levels, undergo information exchange and updating through a multilayer perceptron (MLP) network as follows:

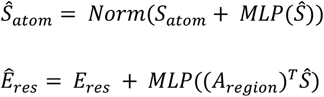

Through iterative updates of residue-level and atomic-level representations, the final enzyme and substrate embeddings are derived using a max-pooling operation. These embeddings are then projected into scalar probability values, facilitating spatially consistent turnover number predictions:

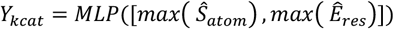

Given the experimental *K*_*cat*_ annotations and the predicted *K*_*cat*_ values, we minimized the root mean squared error (RMSE) loss to optimize the supervised objective. Additionally, we employed the MinCUT loss to learn an unsupervised partitioning of residues, resulting in the following overall loss function:

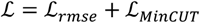

### Experimental materials

#### Plasmid construction

The gene encoding GH13_31 α-Glucosidase (BspAG13_31A) from *Bacillus sp* AHU2216 [51] was codon-optimized, synthesized and cloned into the pET28a vector by Beijing Tsingke Biotech Co. Ltd., resulting in the plasmid pET28a-31A (see Supporting materials). The BspAG13_31A mutants (D69A, H89K, R230Q, W287F, D396K, T408N, Q468D, H528A, H544N) were generated using whole plasmid PCR with the 2x Phanta Max Mix (Vazyme, Nanjing) and pET28a-31A as the template. The primers used are listed in Table S1. The PCR products were digested with DpnI and transformed into *Escherichia coli* DH5a. Correct mutants were verified by sequencing and subsequently transformed into *E. coli* BL21(DE3) for protein expression.

#### Protein expression and purification

The mutants were individually inoculated into 2 mL of LB medium containing 50 μg/mL kanamycin and cultured overnight at 37 °C and 220 rpm. The overnight culture (1 mL) was transferred into 100 mL of LB medium with 50 μg/mL kanamycin in 100 mL shaking flasks. The cultures were incubated at 37 °C and 220 rpm for 2 - 3 h until the OD600 reached 0.6-0.8. Isopropyl β-D-1-thiogalactopyranoside (IPTG) was then added to a final concentration of 0.2 mM, and the cultures were further incubated at 18 °C overnight. After induced expression, the cells were harvested by centrifugation at 4,000 rpm nd 4 °C for 10 minutes. The cell pellets were resuspended in 10 mM phosphate buffer (pH 7.0) containing 0.5 M NaCl and disrupted by sonication. The lysate was centrifuged at 12,000 rpm and 4 °C for 10 minutes, and the supernatant was collected and loaded onto a nickel affinity column (GE Healthcare) for purification. The proteins were desalted using a PD-10 Desalting Column (Cytiva) and analyzed by SDS-PAGE. Protein concentrations were determined using the Bradford Protein Assay Kit (Beyotime, China).

#### Determination of kinetic parameters

Enzymatic assays to determine kinetic parameters were performed in 50 mM NaH2PO4-Na2HPO4 buffer (pH 7.0) at 37 °C using p-nitrophenyl-α-D-glucopyranoside (pNPG) at concentrations ranging from 0 to 5 mM (0 mM, 0.5 mM, 1 mM, 1.25 mM, 2 mM, 2.5 mM, 3.75 mM, 5 mM). Enzyme solutions (0.01∼0.02 μM) were used for different variants. Following the literature [51], reactions were incubated at 37 °C for 15 minutes and then terminated with an equal volume of 100 mM NaCO3. Each reaction was performed in duplicate. Absorbance values were measured at 400 nm, and a standard curve was established to correlate p-nitrophenol (pNP) concentration with optical density (OD400) (Fig. S18). Initial reaction velocities (V0) were calculated from the rate of pNP production per second due to enzymatic hydrolysis at different pNPG concentrations. The V0 values and their corresponding pNPG concentrations were imported into the GraphPad Prism software, which automatically calculated the kinetic parameters (*V*_*max*_, *K*_*m*_, and *K*_*cat*_) using the Michaelis-Menten equation.

## Acknowledgements

We thank all providers of public data used in this work for their sharing. In particular, we thank Robin B. Gasser for valuable insights during the manuscript revision.

## Author’s contributions

T.P. and J.S. conceived the ideas. T.P., X.C. and G.Z. designed the experiments. T.P. and X.C. performed the experiments. T.P. analysed the data and prepared figures. T.P. and J.S. wrote the manuscript. G.I.W. and L.K. provided guidance on data analyses. G.Z., G.I.W. and J.S. supervised the project. All authors contributed ideas to the work and assisted in manuscript editing and revision.

## Data availability

All raw and benchmark data resources used in this study are publicly available via Zenodo at (https://doi.org/10.5281/zenodo.14997549). The curated datasets were obtained from the following databases: BRENDA (https://www.brenda-enzymes.org/), SABIO-RK (<u>http://sabio.h-its.org/),</u> Protein Data Bank (PDB) (https://www.rcsb.org), UniProt (https://www.uniprot.org), and M-CSA (https://www.ebi.ac.uk/thornton-srv/m-csa/). All source code required to evaluate the conclusions in the paper is provided in the paper and/or the Supplementary Materials and can be accessed at https://github.com/BioColLab/KcatNet.

## Declarations

### Ethics approval and consent to participate

Not applicable.

### Consent for publication

Not applicable.

---

**Extended Data Figure 1.**
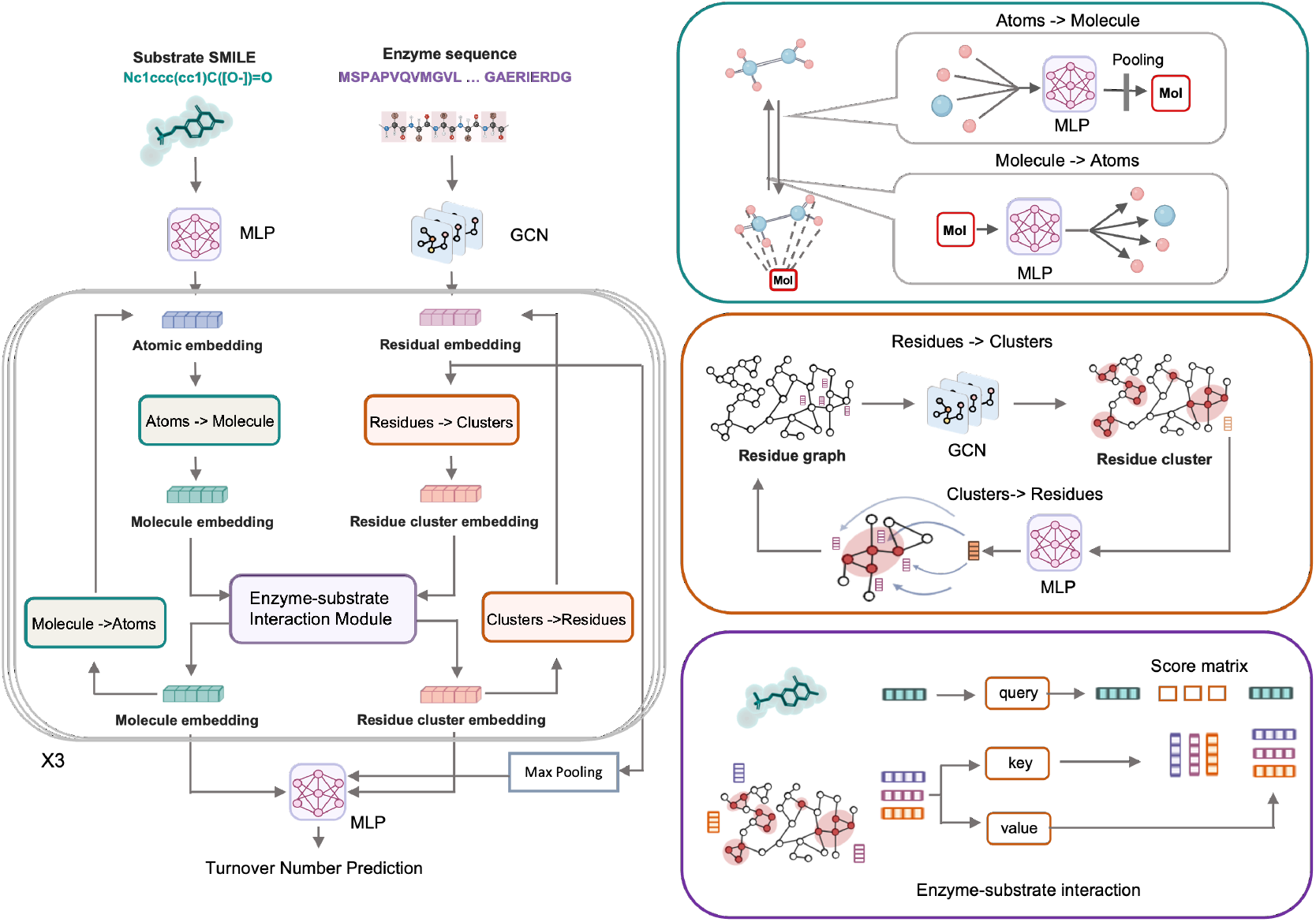
Schematic diagram of KcatNet architecture for predicting enzyme turnover numbers. The substrate representations are processed through a Multi-Layer Perceptron (MLP) to update atomic embeddings, while the enzyme representations are passed through a graph convolutional network (GCN) to obtain updated residue embeddings. These embeddings are further refined through the Atoms-to-Molecule and Residues-to-Clusters modules to generate molecule and residue cluster embeddings, respectively. The enzyme-substrate interaction module facilitates message passing between the substrate and residue clusters, refining the molecule and residue cluster embeddings. Additional transformations, including Molecule-to-Atom and Residue Cluster-to-Residue mappings, are applied before the embeddings are fed into a final MLP for turnover number prediction, with max pooling applied to the residue-level embeddings. Substrate representation is embedded at both atomic and molecular levels, with the Atoms-to-Molecule module passing messages from atomic to molecular embeddings, weighted by importance scores calculated through an MLP. The Molecule-to-Atom module allows the reverse process. The Residues-to-Clusters module groups residues into clusters based on spatial proximity using GCN graph pooling, while the Cluster-to-Residue mapping unpools the clustered regions. The enzyme-substrate interaction module uses cross-attention to capture interactions between enzyme residue clusters and substrates. Refer to the Method section for details.

**Extended Data Figure 2.**
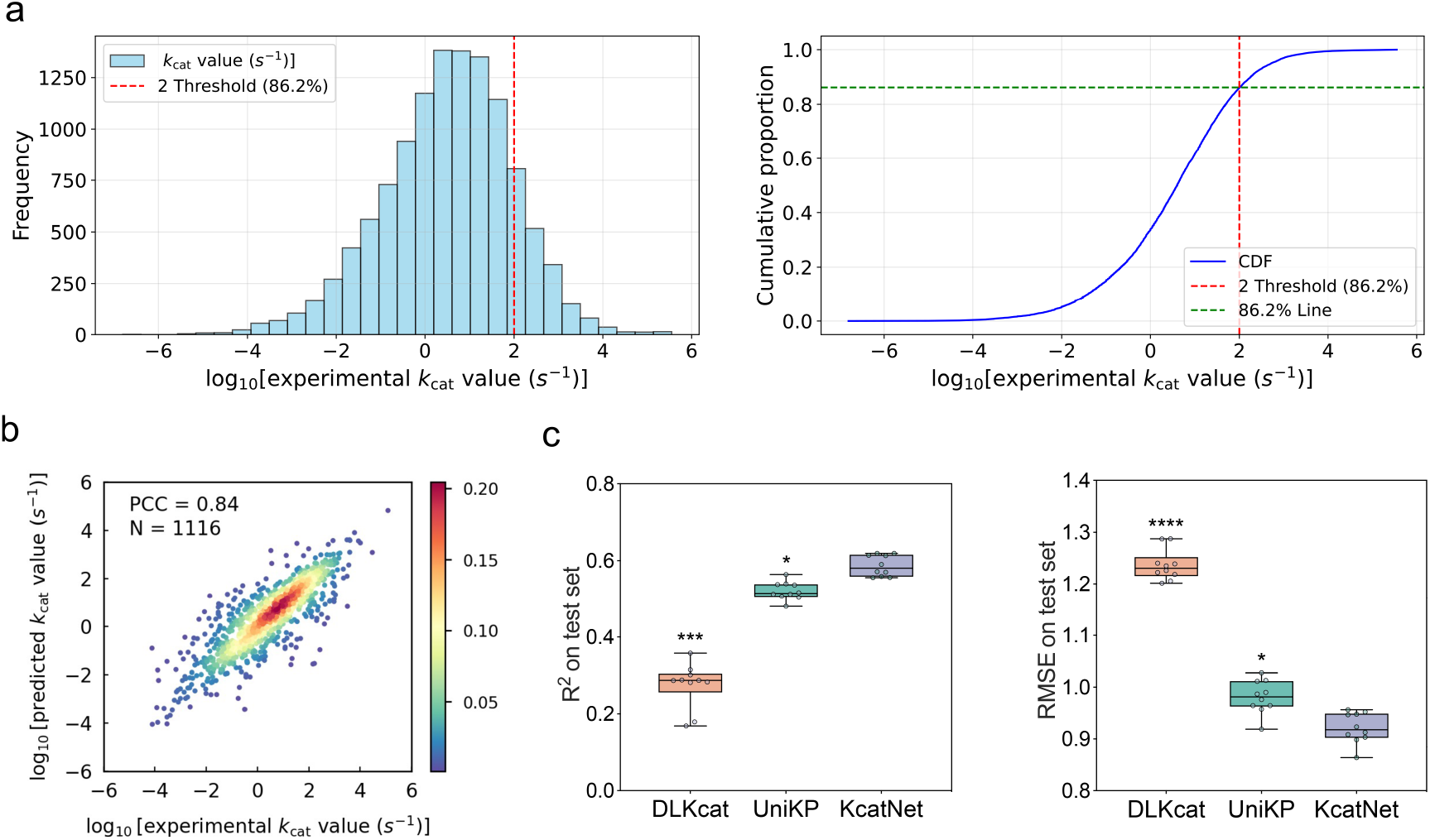
Distribution of *K*_cat_ values and model performance on unseen enzymes. a Distribution of *K*_*cat*_ values in the entire dataset (left). Cumulative distribution function (CDF) of the experimental *K*_*cat*_ values (right). The green dashed line represents the 86.2% threshold for reference, and the red dashed line indicates the *K*_*cat*_ values corresponding to the 86.2% threshold of the data. b Scatter plot showing the Pearson coefficient correlation (PCC) between experimentally measured *K*_*cat*_ values and KcatNet-predicted *K*_*cat*_ values for novel enzymes (***n*** = 1116) in the test dataset. c Comparison of KcatNet with baseline models DLKcat and UniKP based on the Coefficient of Determination (R2) (left) and Root Mean Square Error (RMSE) (right) for novel enzymes in the test dataset. For boxplots, the center line represents the median, while the top and bottom edges mark the first and third quartiles, respectively. Statistical significance levels are indicated (two-sided ***t***-test: ∗, ***P*** ≤ 0.05; ∗∗, ***P*** ≤ 0.01; ∗∗∗, ***P*** ≤ 0.001; ∗∗∗∗, ***P*** ≤ 0.0001).

**Extended Data Figure 3.**
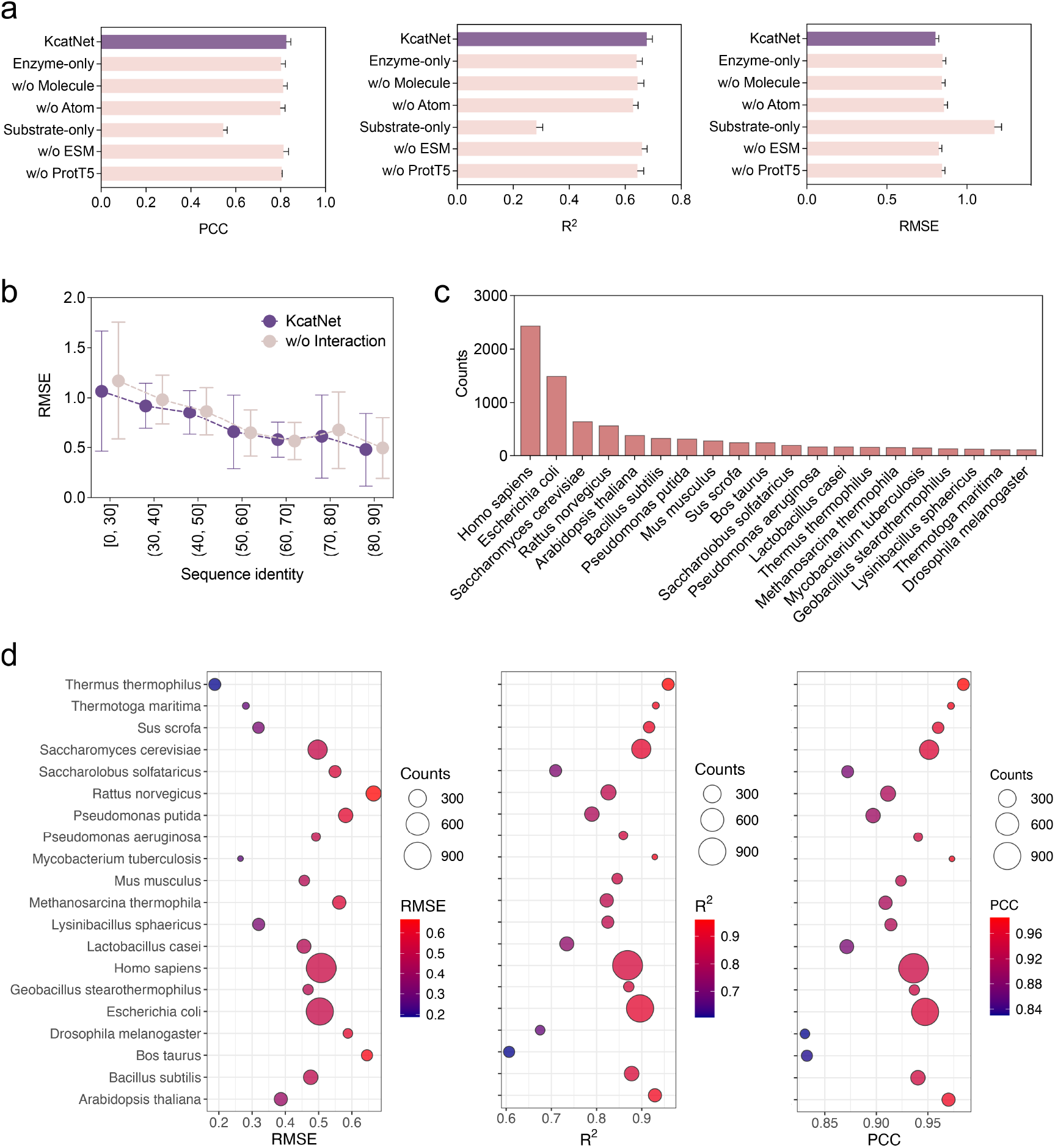
Ablation study of KcatNet and application of KcatNet to enzymes in various organisms. **a** Importance of different input features for KcatNet performance on the test datasets. Feature importance was assessed by removing specific features and evaluating their impact on model performance in terms of PCC (left), R^2^ (medium), and RMSE (right). Considered features included enzyme representation-based features (embeddings from ProtT5 and ESM models) and substrate representation-based features (atom-level features using atom type and property embeddings, and molecule-level features processed via a pretrained SMILES transformer). **b** Comparison of model performance at varying sequence identities, where the enzyme-substrate interaction module of KcatNet is excluded (denoted as w/o Interaction). Instead, substrate and enzyme features are concatenated directly. Excluding the interaction module leads to reduced performance, highlighting the importance its importance in the model. **c** Enzymes in the test dataset are grouped by their organisms, showing the enzyme counts within each organismic category. **d** Performance of KcatNet in predicting *in vitro K*_*cat*_ values for enzymes across diverse organisms, evaluated using RMSE, R^2^. and PCC.

**Extended Data Figure 4.**
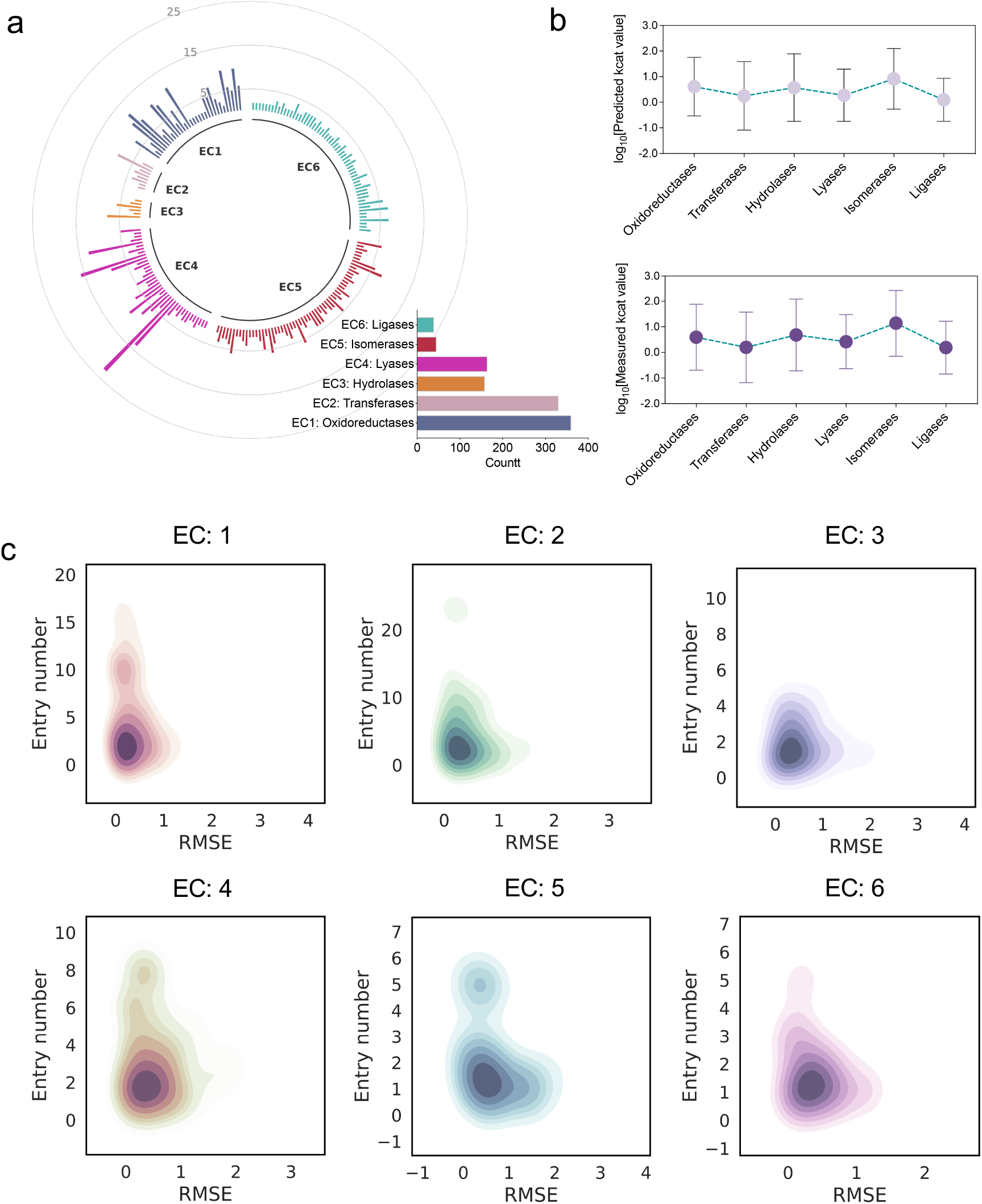
Application of KcatNet to enzymes with varying functions. **a** Primary EC number-based categorization of enzymes: Enzymes in the test dataset are grouped by their primary EC number, showing the enzyme counts within each functional category. **b** Predicted *K*_*cat*_ values by KcatNet for enzymes within each functional category (upper panel) and experimental *K*_*cat*_ values (lower panel) for enzymes across all EC classes. **c** Effect of the number of identical catalytic reactions in the training dataset on predictive performance across enzyme function categories.

## References

1. Markin C, Mokhtari D, Sunden F, Appel M, Akiva E, Longwell S, Sabatti C, Herschlag D, Fordyce P: Revealing enzyme functional architecture via high-throughput microfluidic enzyme kinetics. Science 2021, 373(6553):eabf8761.

2. Basan M, Hui S, Okano H, Zhang Z, Shen Y, Williamson JR, Hwa T: Overflow metabolism in Escherichia coli results from efficient proteome allocation. Nature 2015, 528(7580):99–104.

3. Zhu J, Thompson CB: Metabolic regulation of cell growth and proliferation. Nature reviews Molecular cell biology 2019, 20(7):436–450.

4. Nguyen V, Wilson C, Hoemberger M, Stiller JB, Agafonov RV, Kutter S, English J, Theobald DL, Kern D: Evolutionary drivers of thermoadaptation in enzyme catalysis. Science 2017, 355(6322):289–294.

5. Domenzain I, Sánchez B, Anton M, Kerkhoven EJ, Millán-Oropeza A, Henry C, Siewers V, Morrissey JP, Sonnenschein N, Nielsen J: Reconstruction of a catalogue of genome-scale metabolic models with enzymatic constraints using GECKO 2.0. Nat Commun 2022, 13(1):3766.

6. Fang X, Lloyd CJ, Palsson BO: Reconstructing organisms in silico: genome-scale models and their emerging applications. Nat Rev Microbiol 2020, 18(12):731–743.

7. Campbell E, Kaltenbach M, Correy GJ, Carr PD, Porebski BT, Livingstone EK, Afriat-Jurnou L, Buckle AM, Weik M, Hollfelder F: The role of protein dynamics in the evolution of new enzyme function. Nat Chem Biol 2016, 12(11):944–950.

8. Lalanne J-B, Taggart JC, Guo MS, Herzel L, Schieler A, Li G-W: Evolutionary convergence of pathway-specific enzyme expression stoichiometry. Cell 2018, 173(3):749-761. e738.

9. Eisenmesser EZ, Millet O, Labeikovsky W, Korzhnev DM, Wolf-Watz M, Bosco DA, Skalicky JJ, Kay LE, Kern D: Intrinsic dynamics of an enzyme underlies catalysis. Nature 2005, 438(7064):117–121.

10. Chen K, Arnold FH: Engineering new catalytic activities in enzymes. Nat Catal 2020, 3(3):203–213.

11. Wendering P, Arend M, Razaghi-Moghadam Z, Nikoloski Z: Data integration across conditions improves turnover number estimates and metabolic predictions. Nat Commun 2023, 14(1):1485.

12. Park JO, Rubin SA, Xu Y-F, Amador-Noguez D, Fan J, Shlomi T, Rabinowitz JD: Metabolite concentrations, fluxes and free energies imply efficient enzyme usage. Nat Chem Biol 2016, 12(7):482–489.

13. Schomburg I, Chang A, Schomburg D: BRENDA, enzyme data and metabolic information. Nucleic Acids Res 2002, 30(1):47–49.

14. Wittig U, Kania R, Golebiewski M, Rey M, Shi L, Jong L, Algaa E, Weidemann A, Sauer-Danzwith H, Mir S: SABIO-RK—database for biochemical reaction kinetics. Nucleic Acids Res 2012, 40(D1):D790-D796.

15. Heckmann D, Lloyd CJ, Mih N, Ha Y, Zielinski DC, Haiman ZB, Desouki AA, Lercher MJ, Palsson BO: Machine learning applied to enzyme turnover numbers reveals protein structural correlates and improves metabolic models. Nat Commun 2018, 9(1):5252.

16. Adams JA: Kinetic and catalytic mechanisms of protein kinases. Chemical reviews 2001, 101(8):2271–2290.

17. Otten R, Liu L, Kenner LR, Clarkson MW, Mavor D, Tawfik DS, Kern D, Fraser JS: Rescue of conformational dynamics in enzyme catalysis by directed evolution. Nat Commun 2018, 9(1):1314.

18. Planas-Iglesias J, Marques SM, Pinto GP, Musil M, Stourac J, Damborsky J, Bednar D: Computational design of enzymes for biotechnological applications. Biotechnol Adv 2021, 47:107696.

19. Li F, Yuan L, Lu H, Li G, Chen Y, Engqvist MK, Kerkhoven EJ, Nielsen J: Deep learning-based k cat prediction enables improved enzyme-constrained model reconstruction. Nat Catal 2022, 5(8):662–672.

20. Kroll A, Rousset Y, Hu X-P, Liebrand NA, Lercher MJ: Turnover number predictions for kinetically uncharacterized enzymes using machine and deep learning. Nat Commun 2023, 14(1):4139.

21. Yu H, Deng H, He J, Keasling JD, Luo X: UniKP: a unified framework for the prediction of enzyme kinetic parameters. Nat Commun 2023, 14(1):8211.

22. Bentéjac C, Csörgő A, Martínez-Muñoz G: A comparative analysis of gradient boosting algorithms. Artificial Intelligence Review 2021, 54:1937–1967.

23. Geurts P, Ernst D, Wehenkel L: Extremely randomized trees. Machine learning 2006, 63:3–42.

24. Koh HY, Nguyen AT, Pan S, May LT, Webb GI: Physicochemical graph neural network for learning protein–ligand interaction fingerprints from sequence data. Nature Machine Intelligence 2024:1-15.

25. Mitra R, Li J, Sagendorf JM, Jiang Y, Cohen AS, Chiu T-P, Glasscock CJ, Rohs R: Geometric deep learning of protein–DNA binding specificity. Nat Methods 2024, 21(9):1674–1683.

26. Choudhury S, Narayanan B, Moret M, Hatzimanikatis V, Miskovic L: Generative machine learning produces kinetic models that accurately characterize intracellular metabolic states. Nat Catal 2024, 7(10):1086–1098.

27. Lin Z, Akin H, Rao R, Hie B, Zhu Z, Lu W, Smetanin N, Verkuil R, Kabeli O, Shmueli Y: Evolutionary-scale prediction of atomic-level protein structure with a language model. Science 2023, 379(6637):1123–1130.

28. Leveson-Gower RB, Mayer C, Roelfes G: The importance of catalytic promiscuity for enzyme design and evolution. Nature Reviews Chemistry 2019, 3(12):687–705.

29. Pan T, Bi Y, Wang X, Zhang Y, Webb GI, Gasser RB, Kurgan L, Song J: SCREEN: A Graph-based Contrastive Learning Tool to Infer Catalytic Residues and Assess Enzyme Mutations. Genomics Proteomics Bioinformatics 2025:qzae094.

30. Falivene L, Cao Z, Petta A, Serra L, Poater A, Oliva R, Scarano V, Cavallo L: Towards the online computer-aided design of catalytic pockets. Nat Chem 2019, 11(10):872–879.

31. Elnaggar A, Heinzinger M, Dallago C, Rihawi G, Wang Y, Jones L, Gibbs T, Feher T, Angerer C, Steinegger M: ProtTrans: Towards cracking the language of Life’s code through self-supervised deep learning and high performance computing. arXiv 2020. arXiv preprint arXiv:200706225 2007.

32. Honda S, Shi S, Ueda HR: Smiles transformer: Pre-trained molecular fingerprint for low data drug discovery. arXiv preprint arXiv:191104738 2019.

33. Corso G, Cavalleri L, Beaini D, Liò P, Veličković P: Principal neighbourhood aggregation for graph nets. Advances in Neural Information Processing Systems 2020, 33:13260–13271.

34. Jia Z, Lin S, Gao M, Zaharia M, Aiken A: Improving the accuracy, scalability, and performance of graph neural networks with roc. Proceedings of Machine Learning and Systems 2020, 2:187–198.

35. Elia I, Haigis MC: Metabolites and the tumour microenvironment: from cellular mechanisms to systemic metabolism. Nature metabolism 2021, 3(1):21–32.

36. Das S, Lee D, Sillitoe I, Dawson NL, Lees JG, Orengo CA: Functional classification of CATH superfamilies: a domain-based approach for protein function annotation. Bioinformatics 2015, 31(21):3460–3467.

37. Zhao S, Kumar R, Sakai A, Vetting MW, Wood BM, Brown S, Bonanno JB, Hillerich BS, Seidel RD, Babbitt PC: Discovery of new enzymes and metabolic pathways by using structure and genome context. Nature 2013, 502(7473):698–702.

38. Ribeiro AJM, Holliday GL, Furnham N, Tyzack JD, Ferris K, Thornton JM: Mechanism and Catalytic Site Atlas (M-CSA): a database of enzyme reaction mechanisms and active sites. Nucleic Acids Res 2018, 46(D1):D618-D623.

39. Hegeman AD, Gross JW, Frey PA: Probing catalysis by Escherichia coli dTDP-glucose-4, 6-dehydratase: identification and preliminary characterization of functional amino acid residues at the active site. Biochemistry 2001, 40(22):6598–6610.

40. Murkin AS, Birck MR, Rinaldo-Matthis A, Shi W, Taylor EA, Schramm VL: Neighboring group participation in the transition state of human purine nucleoside phosphorylase. Biochemistry 2007, 46(17):5038–5049.

41. Gerasimavicius L, Livesey BJ, Marsh JA: Loss-of-function, gain-of-function and dominant-negative mutations have profoundly different effects on protein structure. Nat Commun 2022, 13(1):3895.

42. Li G, Hu Y, Zrimec J, Luo H, Wang H, Zelezniak A, Ji B, Nielsen J: Bayesian genome scale modelling identifies thermal determinants of yeast metabolism. Nat Commun 2021, 12(1):190.

43. Chen Y, Nielsen J: In vitro turnover numbers do not reflect in vivo activities of yeast enzymes. Proceedings of the National Academy of Sciences 2021, 118(32):e2108391118.

44. Han G-S, Carman GM: Characterization of the human LPIN1-encoded phosphatidate phosphatase isoforms. J Biol Chem 2010, 285(19):14628–14638.

45. Auiewiriyanukul W, Saburi W, Kato K, Yao M, Mori H: Function and structure of GH 13_31 α-glucosidase with high α-(1→ 4)-glucosidic linkage specificity and transglucosylation activity. FEBS letters 2018, 592(13):2268–2281.

46. Meier J, Rao R, Verkuil R, Liu J, Sercu T, Rives A: Language models enable zero-shot prediction of the effects of mutations on protein function. Advances in neural information processing systems 2021, 34:29287–29303.

47. Dauparas J, Anishchenko I, Bennett N, Bai H, Ragotte RJ, Milles LF, Wicky BI, Courbet A, de Haas RJ, Bethel N: Robust deep learning–based protein sequence design using ProteinMPNN. Science 2022, 378(6615):49–56.

48. Johnson SR, Fu X, Viknander S, Goldin C, Monaco S, Zelezniak A, Yang KK: Computational scoring and experimental evaluation of enzymes generated by neural networks. Nat Biotechnol 2024:1–10.

49. Cocco S, Posani L, Monasson R: Functional effects of mutations in proteins can be predicted and interpreted by guided selection of sequence covariation information. Proceedings of the National Academy of Sciences 2024, 121(26):e2312335121.

50. Mader MM, Bartlett PA: Binding energy and catalysis: the implications for transition-state analogs and catalytic antibodies. Chemical reviews 1997, 97(5):1281–1302.

51. Kobayashi M, Hondoh H, Mori H, Saburi W, Okuyama M, Kimura A: Calcium ion-dependent increase in thermostability of dextran glucosidase from Streptococcus mutans. Bioscience, biotechnology, and biochemistry 2011, 75(8):1557–1563.

